# Universal salt iodization contributes to health equity: socio-economic status does not affect iodine status in children in Macedonia

**DOI:** 10.1101/2020.12.24.424286

**Authors:** Neda Milevska Kostova, Borislav Karanfilski, Daniela Miladinova, Sonja Kuzmanovska, Biljana Chuleva, Venjamin Majstorov, Till Ittermann, Henry Völzke

## Abstract

Many studies have shown that socio-economic status (SES) contributes to health inequalities, with nutrition as one of the major risk factors. Iodine intake entirely depends on external sources, and deficiencies are known to be more prevalent in lower social groups, especially in countries with limited access to iodized salt. This study aimed to determine the influence of SES on iodine status and iodine availability from household salt in North Macedonia. Using cluster sampling, 1,200 children were recruited, and 1,191 children participated (response rate: 99.2%). Iodine status was assessed through urinary iodine concentration (UIC), and iodine availability through iodine content in household salt requested from participants. SES was assessed using standardized Family Affluence Score (FAS). No statistically significant correlation was found between FAS and iodine in salt. Median regression revealed no significant associations of middle vs. low FAS (β=0.00; 95%-confidence interval (CI)=[−0.61, 0.62]; p=0.999) or high vs. low affluence (β=0.48; 95%CI=[−1.37, 0.41]; p=0.291) with iodine content in household salt. UIC levels were significantly lower in middle FAS children compared to low FAS children (β=-16.4; 95%CI=[−32.3,-0.5]; p=0.043). No statistically significant differences in UIC were found between children with high and low affluence (β=-12.5; 95%CI=[−35.5, 10.5]; p=0.287), possibly due to lowered statistical power for this comparison. Universal salt iodization (USI) proves to be cost-effective measure for appropriate iodine intake in healthy children and adults, irrespective of their social status. It can thus be concluded that USI contributes to reducing health inequalities related to iodine status among population of different social strata.

## Introduction

The association between socio-economic status (SES) and proper nutrition is well documented (1, 2). One key element of diet quality is the micronutrient content (3), and recent reviews showed that the prevalence of micronutrient inadequacies in Europe ranged from 11% to 30% for copper, folate, selenium, vitamin B12, vitamin C and iodine in adults (3, 4). Iodine intake entirely depends on its availability in food, and its deficiency has been identified as one of the modifiable factors that have an adverse effect on child development (5). Iodine deficiencies are known to be more prevalent in lower social groups, especially in countries with limited access to iodized salt (6).

Despite substantial progress made, iodine deficiency remains a significant health problem worldwide and affects both developed and developing nations (7). Studies on iodine intake for children in Europe reported lower intake in low-SES than in high-SES groups (8, 9). However, such studies for Central and Eastern Europe are very limited (3).

To ensure adequate iodine availability and intake, in North Macedonia in 1956, initiative for mandatory iodization of household salt with 10 mg of potassium iodide (KI) per kg salt was raised, but high prevalence of goitre persisted (10). In 1999, the USI policy was changed to iodization of entire edible salt with 20-30 mg of a more stable potassium iodate (KIO_3_) per kg salt, including both for household use and food industry. The results of such efforts have shown reduction of goitre prevalence from 18.7% in 1996 (before the new USI policy) to 4.7% in 2003 (11). The median urinary iodine concentration (UIC), from 117μg/L in 1996 increased to 191μg/L in 2003 (11).

In none of the previous Macedonian studies SES has been taken into consideration. The objective of this study is to determine the effects of USI on reducing health inequalities, through assessing the influence of SES on iodine status of school children and iodine availability from household salt in North Macedonia.

## Materials and Methods

### Study population

The SES survey was undertaken as part of a regular iodine status assessment study performed periodically in the country. Study sample of 1,200 children between 8-10 years of age was chosen using random sampling from the list of all primary schools and number of children enrolled in the 3^rd^ and 4^th^ grade, obtained from the Ministry of Education and Science for the school year 2015/2016. The sample was defined as 30 clusters of 40 children, where cluster size (2,030 children) was determined by dividing the total number of children in the 8-10 years’ cohort with the number of clusters. Schools were chosen from an alphabetical list of schools and cumulative number of enrolled children, to which the cluster size was applied, then choosing the nearest one. School directors were requested to randomly select 20 girls and 20 boys, to inform parents and obtain written informed consent. The sampling and data collection was conducted between March and October 2016. Of the recruited children, 1,191 participated (response 99.2%), of which 589 were girls (49.5%) and 602 were boys (50.5%). The median age was 9 years (mean: 8.6), and median height and weight were 137 cm (mean: 137.2 cm) and 31.0 kg (mean: 33.1 kg), respectively (Table 1).

**Table 1.** Basic Characteristics of examined school children (n=1,191)

| **Characteristics** | **Frequency (person)** | **Percentage (%)** |
| --- | --- | --- |
| **Age** |  |  |
| Up to 8 years old | 66 | 5.5% |
| 8 to 9 years old | 489 | 41.1% |
| 9 to 10 years old | 636 | 53.4% |
| **Gender** |  |  |
| Girls | 589 | 49.5% |
| Boys | 602 | 50.5% |
| **Ethnicity** |  |  |
| Macedonian | 687 | 57.7% |
| Albanian | 429 | 36.0% |
| Turkish | 6 | 0.5% |
| Roma | 60 | 5.0% |
| Other | 9 | 0.8% |
| **Place of residence (according to the school location)** |  |  |
| Urban – capital | 334 | 28.0% |
| Urban – other | 564 | 47.4% |
| Rural | 293 | 24.6% |
| **Distribution of sample by UIC adequacy clusters** |  |  |
| <100 µg/L | 106 | 9.1% |
| 100-199 µg/L | 298 | 25.5% |
| 200-299 µg/L | 447 | 38.3% |
| >300 µg/L | 316 | 27.1% |
| n/a | 24 |  |
| **Family Affluence Score (FAS)** |  |  |
| FAS 1 (score=0–3) low affluence | 385 | 32.3% |
| FAS 2 (score=4,5) middle affluence | 641 | 53.8% |
| FAS 3 (score=6,7) high affluence | 165 | 13.9% |

### Measurements

The study consisted of several measurements, including assessment of iodine status through urinary iodine concentration, iodine availability through household salt and assessment of socio-economic status through questionnaire. All urinary samples were taken between 9 and 15 o’clock on the examination day. Household samples were obtained from the recruited children who were asked to provide about 100 grams of salt currently used in their home, together with producer/importer name and production and expiry dates of the salt, as written on the original package. To avoid misinterpretation of questions, the questionnaire was filled-in by a study team member with responses given by children, at the same time confirming whether they have understood what they were asked.

Iodine content in household salt was determined using iodometric titration method, which gives the exact value of iodine in the salt in ppm (12, 13). UIC was determined using Sandell-Kolthoff reaction, based on the role of iodide as a catalyst in the reduction of ceric ammonium sulphate to the cerous form, in the presence of arsenious acid (14). The absorbance was determined at 420nm using UV/VIS spectrophotometer (UNICO UV-2102, Unico Instrument Co., Ltd., USA). As part of the harmonisation of methods activity within the EUthyroid initiative, the measured UIC values were adjusted using correction formula (UICcorr = 1.04*UIC-29.21) obtained through validation of 75 random samples measured in the EUthyroid reference laboratory THL in Helsinki, Finland, using reference method-Inductively Coupled Plasma Mass Spectrometry (ICP-MS) (15).

SES is usually determined using specific social or economic parameters, such as level of income or education (6). As the survey was conducted in minor children, the possibility of obtaining such information from the sample was rejected for several reasons: 1) the children are likely to not know this information about their family; 2) they might not have understanding of the parity of financial assets in general; and 3) they might not be aware of their right to privacy. Considering that SES assessment was intended as a proxy measure for the impact of social and economic conditions on iodine status, we have used the index of affluence, already used for similar purposes (16, 17). The Family Affluence Score (FAS) measures the socio-economic status indirectly through access to certain commodities or services in everyday life (16, 18). Thus, for assessment of SES, children were asked to respond to a short form of the standardized Health-behaviour of school children (HBSC) survey questionnaire consisting of 4 questions: 1) Does your family own a car, van or truck? 2) Do you have your own bedroom? 3) During the past 12 months, how many times did you travel with your family on holiday? and 4) How many computers does your family own?. The FAS was calculated based on a response matrix, using a three-point ordinal scale (Table 2).

**Table 2.** Guide for statistical processing of SES questionnaire responses.

| Question | (Family Affluence Score) |  |  |  |
| --- | --- | --- | --- | --- |
|  | 0 | 1 | 2 | 3 |
| Does your family own a car, van or truck? | No | Yes, one | Yes, two or more | n/a |
| Do you have your own bedroom for yourself? | No | Yes |  |  |
| During the past 12 months, how many times did you travel away on holiday with your family? | Not at all | Once | Twice | More than twice |
| How many computers does your family own? | None | One | Two | More than two |
| **FAS three-point ordinal scale** |  |  |  |  |
| • FAS 1 (score = 0–3) indicated low affluence<br>• FAS 2 (score = 4, 5) indicated middle affluence<br>• FAS 3 (score = 6, 7) indicated high affluence |  |  |  |  |

The calculated FAS was divided into a scale of three subcategories (low, middle and high affluence) and compared to the values of iodine content in household salt and of UIC to determine the strength of associations between them and whether the differences are statistically significant.

### Statistical analysis

Data on quantitative characteristics are expressed as median and interquartile range. Simple descriptive analyses are used to present the distribution of responses on SES, and values of iodine content in cooking salt and UIC. Correlations were determined using the Spearman rank-order correlation. Furthermore, we performed median regressions to associate the FAS with UIC and iodine content in household salt. The regressions were adjusted for age, sex and place of residence (capital, urban, rural).

A p-value of <0.05 was considered statistically significant. All calculations were performed using Small STATA 15.1 (Stata Corporation, College Station, TX).

## Results

### FAS and salt iodization

Salt iodization levels were determined in 1,092 samples with average iodine content of 24.8 mg/kg (range 0.7-58.2 mg/kg) and median of 24.3 mg/kg (p25=22.2; p75=27). We compared the average levels of salt iodization and average FAS values by school (Figure 1), using Spearman’s rank correlation, which showed no significant correlation (r_s_= 0.0042).

**Figure 1.**
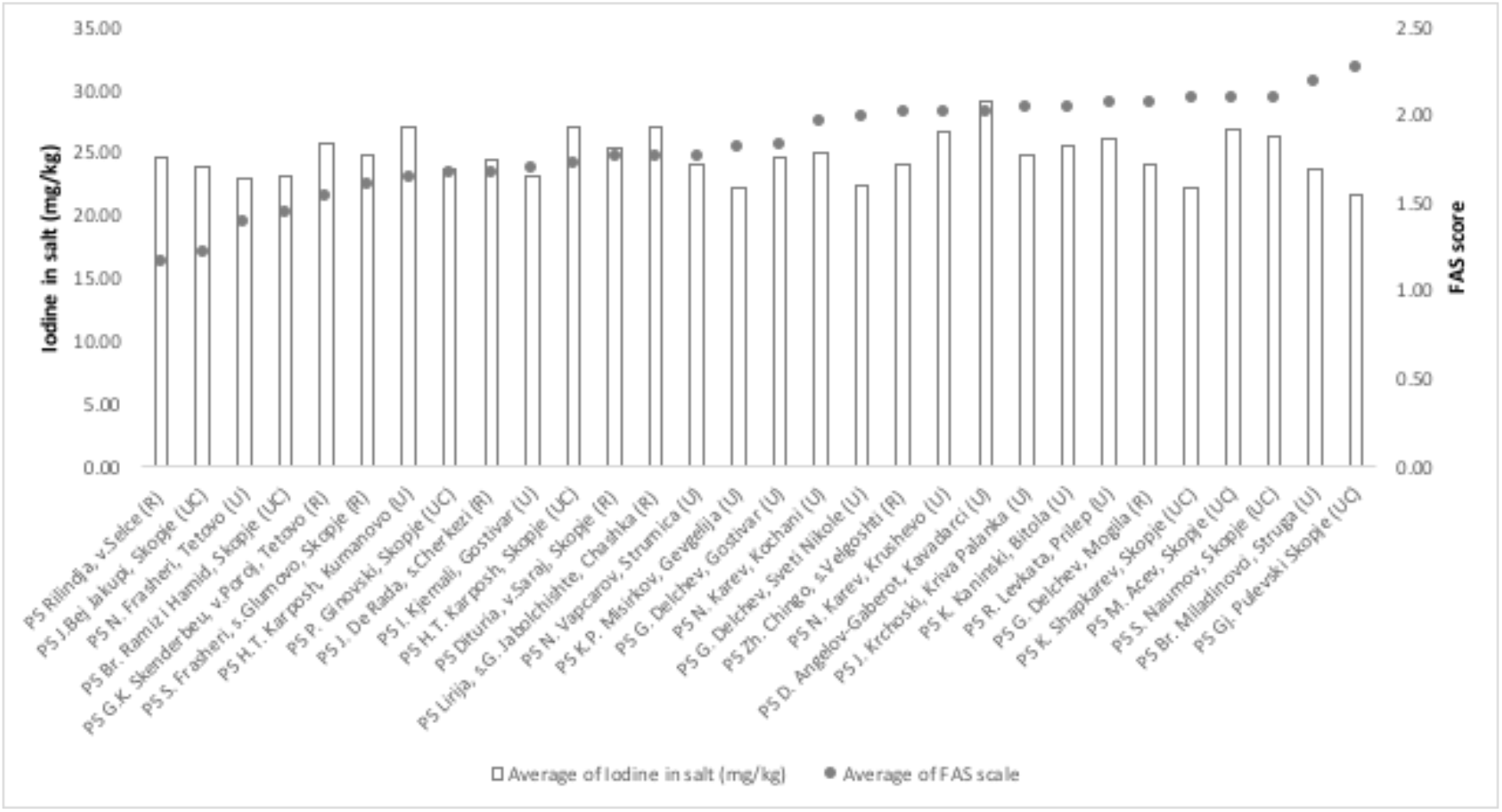
Comparison of average iodine content in household salt and FAS by cluster. Legend: School in: R = rural; U = urban; UC = urban capital area

Multivariable median regression revealed no significant associations of middle vs. low affluence (β=0.00; 95% confidence interval (CI) = −0.61 to 0.62; p = 0.999) or high vs. low affluence (β=0.48; 95% CI = −1.37 to 0.41; p = 0.291) with iodine content in household salt.

### FAS and UIC

UIC was determined in 1,167 samples, and after adjustments with the correction formula, the average UIC was 242.7 μg/L and median value 236 μg/L (p25=171.3 μg/L; p75=307.6 μg/L) (Table 1).

We compared the average UIC and average FAS values stratified by schools (Figure 2) and we observed no significant association between the SES and UIC.

**Figure 2.**
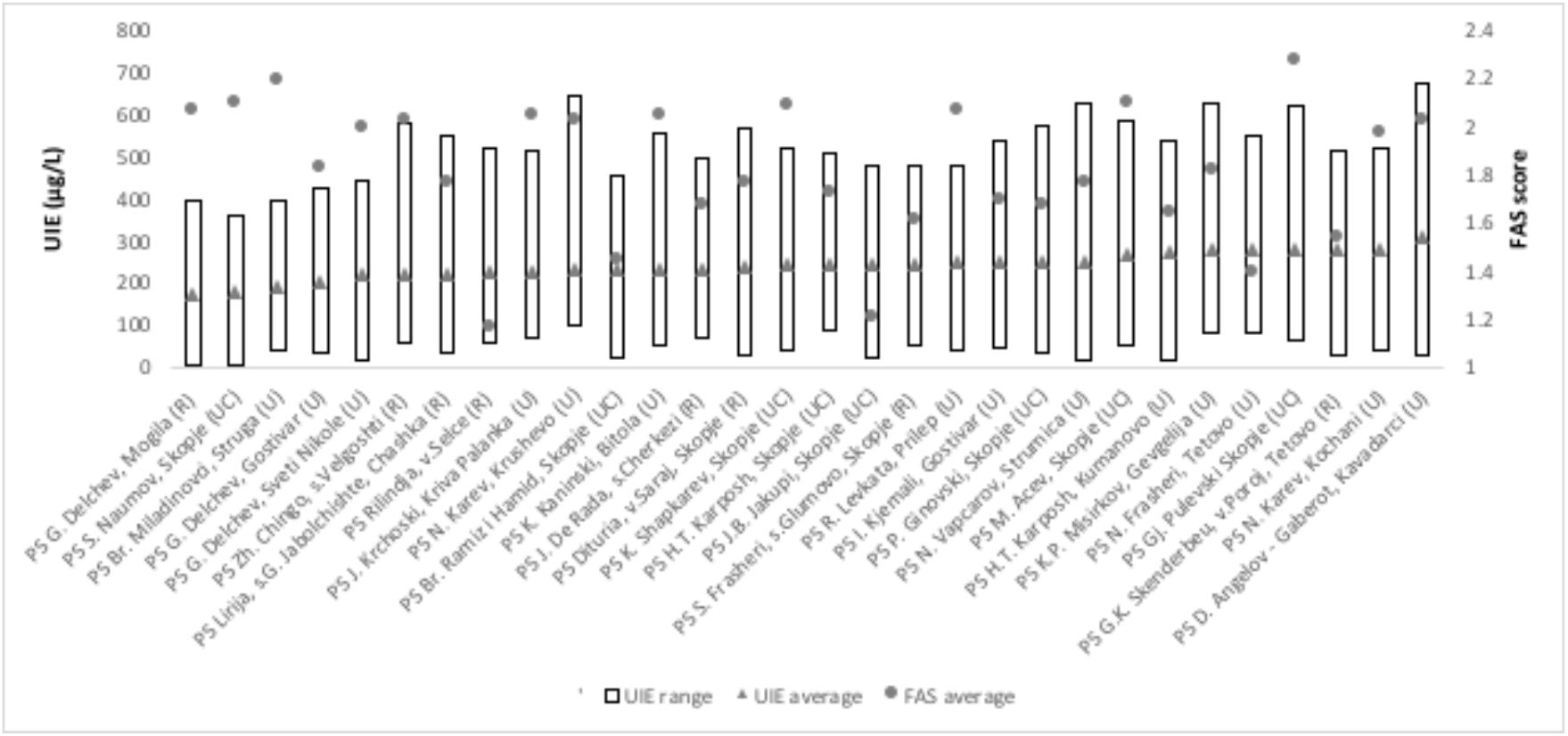
Comparison of UIC range and UIC average with FAS average by cluster. Legend: School in: R = rural; U = urban; UC = urban capital area

Further, using Spearman’s rank correlation, we compared these two parameters for each individual participant, and found no significant correlation (r_s_= − 0.0242). We also compared the average FAS score to the adequacy clusters of UIC, defined as: <100 = insufficient; ≥100-199 = adequate; 200-299 = above requirements; >300 = excessive (Figure 3). No significant association was determined (r_s_ = −0.012).

**Figure 3.**
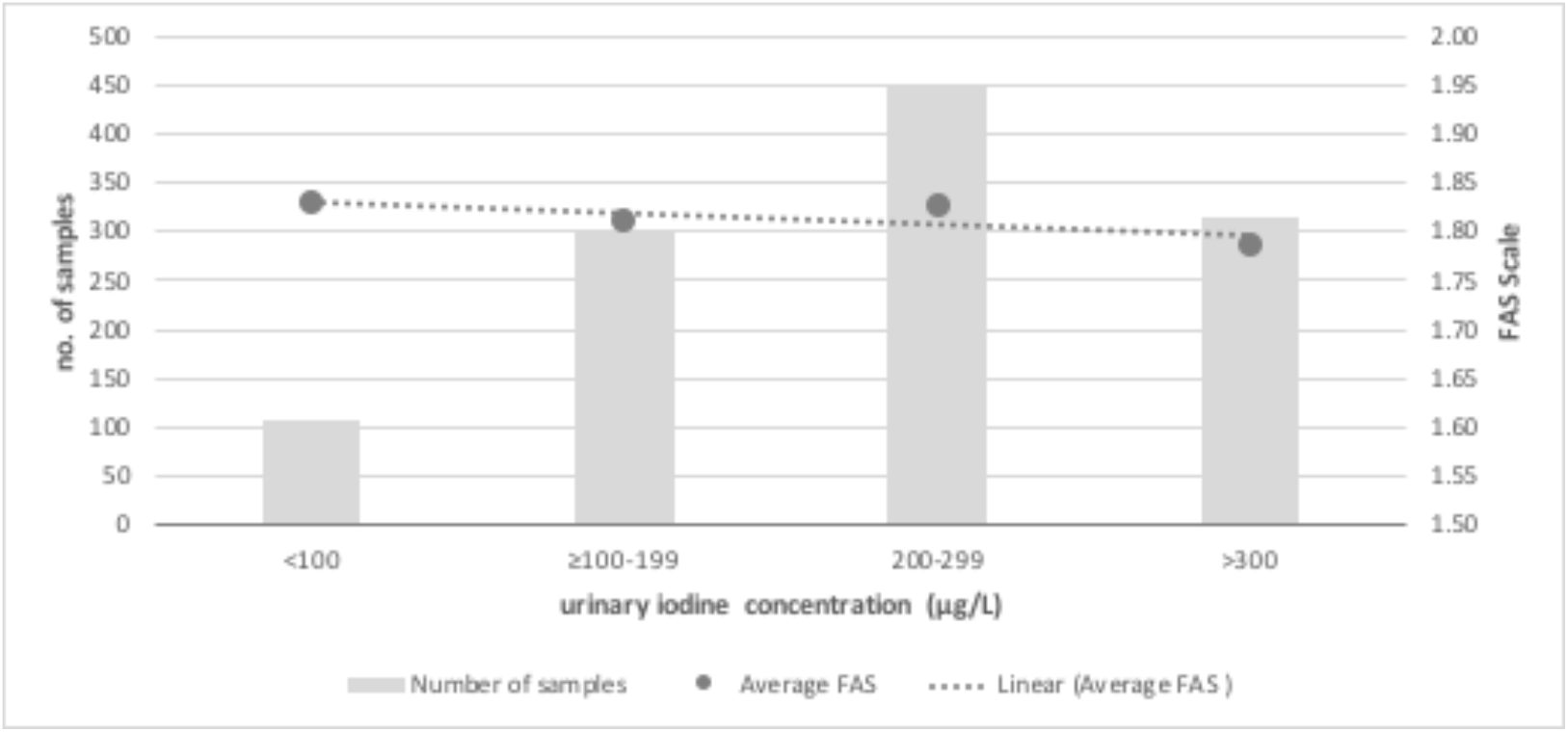
Comparison of average FAS and UIC by adequacy clusters.

In median regression children with a middle affluence score had significantly lower UIC levels compared to children with low affluence score (β=-16.4; 95% CI = −32.3 to −0.5; p = 0.043). No statistically significant differences in the UIC levels were found between children with high and low affluence (β=-12.5; 95% CI = −35.5 to 10.5; p = 0.287).

We also found that the place of residence had no significant effect on UIC levels, but in tendency median UIC levels are highest in the urban and lowest in the rural region (data not shown).

## Discussion

No significant association of SES and iodine content in household salt confirms that the country’s USI policy is properly implemented. More importantly, it suggests that USI policies, in general, ensure health equity through sufficient access to iodine for lower socio-economic groups.

Regarding iodine intake, overall UIC levels lay at the higher end of the recommended values, placing the country in the safe zone for iodine sufficiency. The comparisons have shown no significant association between the SES and UIC, also suggesting sufficient iodine intake in all socio-economic groups. However, the determined association of lower FAS scores with higher UIC, especially between children with a middle and low affluence implies that children of lower SES consume food with higher salt content, which requires further investigation. The absence of such association between children with high and low affluence may be due to the smaller number of children in the high affluence score group, resulting in lowered statistical power for this comparison.

In North Macedonia, the obtained results of the lack of an association between the SES and UIC suggest that USI plays an essential role in ensuring equitable access to iodine for all social classes. This is confirmed with the comparisons of SES and UIC adequacy clusters. Similarly, a study in Northern Germany has not found significant association of socio-economic variables with iodine excretion (19).

European countries lacking specific iodine prophylaxis programs are mostly mildly iodine deficient (20). The prevalence of iodine deficiency in Europe was reduced by 30% from 2003 to 2010, but 44% of school-age children still have insufficient iodine intake (21). In a 2009 survey of schoolgirls aged 14 to 15 years living in the United Kingdom, which has long been considered iodine sufficient, median UIC was 80.1 μg/L, consistent with mild iodine deficiency (22). In Italy, UIC was randomly measured in 26,913 individuals, as part of a project to eradicate iodine deficiency disorders (23). Urinary iodine was lower than 100 and 50 μg/L in 64.3 and 34.9 percent of samples, respectively. Median urinary iodine in nonurban areas was significantly lower than in urban areas (69 μg/L vs. 79 μg/L). However, all of these countries have no programs for universal access to iodine for the entire population, such as universal salt iodization.

The USI policy and efforts of the country in achieving sustainable iodine sufficiency have been acknowledged in a recent review (24). This confirms that universal salt iodization, among other measures, remains one of the most cost-effective measures in elimination of iodine deficiency among children and adult population, given that iodine status of children has been recognized as marker of iodine status of the adult population (25).

## Summary and conclusions

The presented nation-wide iodine status of school children study performed in North Macedonia confirmed adequate iodine supply through table salt and iodine sufficiency measured through UIC. Similar assessments conducted in the past did not take SES into consideration. The added value of this study is that it puts the iodine supply and intake (measured through UIC) in the context of the socio-economic status. A recent review on the subject found lack of such studies, particularly on association between iodine intake and SES in children, concluding on the necessity for further research (3). The random selection of study participants, the nearly complete response and the highly standardized data collections warrant the generalizability of study findings.

This study has shown that USI is a good measure for prevention of iodine deficiency-related disorders of all social groups. However, the finding that lower social strata exhibit higher values of UIC might require further investigation. Also, as the country committed to implementing the WHO recommendations for reducing salt intake, reassessment of iodine status, with particular attention to socio-economic variables is a foreseeable activity in the future.

